# SQANTI-SIM: a simulator of controlled transcript novelty for lrRNA-seq benchmark

**DOI:** 10.1101/2023.08.23.554392

**Authors:** Jorge Mestre-Tomás, Tianyuan Liu, Francisco Pardo-Palacios, Ana Conesa

## Abstract

Long-read RNA-seq has emerged as a powerful tool for transcript discovery, even in well-annotated organisms. However, assessing the accuracy of different methods in identifying annotated and novel transcripts remains a challenge. Here, we present SQANTI-SIM, a versatile utility that wraps around popular long-read simulators to allow precise management of transcript novelty based on the structural categories defined by SQANTI3. By selectively excluding specific transcripts from the reference dataset, SQANTI-SIM effectively emulates scenarios involving unannotated transcripts. Furthermore, the tool provides customizable features and supports the simulation of additional types of data, representing the first multi-omics simulation tool for the lrRNA-seq field. We demonstrate the effectiveness of SQANTI-SIM by benchmarking five transcriptome reconstruction pipelines using the simulated data.

## 1 Introduction

Third-generation or long-read sequencing (TGS/LRS) technologies such as Pacific Biosciences (PacBio) and Oxford Nanopore (ONT) have revolutionized the field of transcriptomics by providing full-length transcripts through single-molecule sequencing spanning kilobase-long reads [1–4]. Transcriptome analysis using long reads (lrRNA-seq) offers several advantages over short-read sequencing, including the identification of alternative isoforms and the analysis of allele-specific expression, and has led to the discovery of numerous novel transcripts even in well-annotated organisms [5, 6]. However, despite recent improvements in TGS technologies, the quality of lrRNA-seq data is compromised by the presence of RNA degradation, library preparation biases, sequencing errors, and mapping artifacts that may result in false transcript models being called from these data. Presently, one of the major challenges in lrRNA-seq data analysis is accurately identifying novel transcripts and distinguishing them from technology artifacts.

Several software tools have been developed for transcript model reconstruction from long reads, which differ, among other features, in their capacity for calling new transcripts. Methods such as TAMA [7], Bambu [8], IsoQuant [9], FLAIR [10] and TALON [11] use the reference annotation to guide transcript calls and are constrained by the existing information. Other approaches such as IsoSeq [12] and LyRic [13] are reference-free and may identify a large number of novel transcripts. Finally, strategies such as StringTie2 [14, 15] assemble together short and long reads resulting in transcript models that may not be fully supported by the long reads. A number of studies have benchmarked lrRNA-seq technologies and tools for accurate isoform identification [2, 7, 16–19] using different combinations of experimental and *in-silico* datasets. Among them, the Long-read RNA-Seq Genome Annotation Assessment Project (LRGASP), a community-oriented assessment [17], stands as the most comprehensive effort to date for evaluating lrRNA-seq methods.

LRGASP included three sequencing platforms, four library preparation methods and fourteen software tools, and used SQANTI3 [20] as quality control tool to guide evaluations of over 50 competing analysis pipelines [17]. SQANTI3 combines annotations and orthogonal data (i.e. Illumina, CAGE and Quant-seq) to evaluate transcript models and proposes a LRS transcript classification scheme based on their comparison to the reference transcriptome. Hence, Full-splice-match (FSM) are LRS transcripts that match a reference transcript at all splice junctions. Incomplete-splice-match (ISM) transcripts miss one or more junctions at 3′ or 5′ end positions, representing either RNA degradation or alternative processing. Novel-in-catalog (NIC) are transcripts with novel combinations of splice sites, while Novel-not-in-catalog (NNC) represent transcript models with at least one novel donor or acceptor site. Other categories are antisense, fusion, intergenic and genic-genomic [21]. Using this scheme the LRGASP project found large discrepancies among lrRNA-seq methods. Especially, the number and identity of transcripts in the novel SQANTI3 structural categories (other than FSM), as well as the level of support by orthogonal data, greatly differed. Remarkably, LRGASP results revealed that many NIC transcripts and a non-neglectable number of NNC, including those identified by a few pipelines, could be validated by targeted PCR amplification [17]. Moreover, several lrRNA-seq studies show that many novel combinations of splice junctions, Transcription Start (TSS), and Termination Sites (TTS), as well as alternative TSS and TTS (i.e. ISM and NIC categories), are likely to be present in RNA samples [18, 20]. These results reflect the grand challenge of the novel transcript discovery from long reads.

Ground-truth strategies used to benchmark RNA-seq include the utilization of spike-ins, experimental validation, and in-silico simulation. Spike-ins or sequins are known synthetic RNA molecules added during the library preparation step that can easily evaluate experimental, sequencing, and mapping biases. However, these kits contain a reduced set of transcripts and have limitations for assessing the complexity of the novel discoveries. Similarly, experimental validation, although highly informative, is costly in time and resources. Data simulation, where the ground truth is decided by the user, is a cost-effective alternative [22, 23]. Simulation algorithms have been developed for PacBio and ONT long-read genomic sequencing [23–26], and also methods specifically designed to simulate lrRNA-seq data are available. Existing tools such as NanoSim [27], IsoSeqSim [28], and PBSIM3 [29] simulate ONT and PacBio transcriptome reads from a reference transcriptome. Unfortunately, these methods do not incorporate a mechanism to simulate novel transcripts, limiting their utility to fully benchmark the lrRNA-seq technology. A few studies have attempted to incorporate novel transcript simulation by introducing other-species transcripts in the reference annotation [17] or eliminating complete chromosomes from the reference GTF when reconstructing transcript models [8]. However, these approaches do not take into account the different types of novel transcripts that arise when evaluating lrRNA-seq data and therefore fail to reveal how experimental and computational algorithms deal with the accurate identification of novel TTS, TSS, splice junctions, and their combinations. Moreover, none of the existing data simulation approaches is able to jointly simulate orthogonal datasets that might be used when defining LRS transcript models, a strategy adopted by some popular lrRNA-seq algorithms [10, 14] and recommended by LRGASP [17]. Therefore the current lrRNA-seq data simulation landscape is insufficient to simulate the complexity and possibilities of the LRS transcriptome data.

Here, we present SQANTI-SIM, an open-source tool designed to enable PacBio and ONT long-read transcriptome data simulation with precise control over the presence of novel transcripts featuring diverse novelty types. Basically, SQANTI-SIM evaluates a user-provided reference GTF to identify transcripts that, when removed from the annotation, become one of the novel SQANTI3 categories. Building on the capabilities of NanoSim, PBSIM3, and IsoSeqSim to simulate long reads from the provided GTF, SQANTI-SIM returns simulated reads and a reduced GTF depleted of transcripts simulated to be novel. Moreover, SQANTI-SIM creates matching Illumina and CAGE data for the synthetic LRS dataset capturing the noisy relationships between data types. By faithfully reproducing LRS transcriptome datasets, SQANTI-SIM provides researchers with a powerful tool for assessing the ability of long-read transcript reconstruction methods to accurately detect known and new transcripts. We apply SQANTI-SIM to different lrRNA-seq algorithms to demonstrate the utility of our tool to reveal distinct abilities for detecting different types of true novel transcripts.

## 2 Results

### 2.1 Overview of SQANTI-SIM workflow

SQANTI-SIM is a lrRNA-seq simulation environment that generates Nanopore dRNA and cDNA reads, as well as PacBio cDNA reads with precise control of transcript novelty based on SQANTI3 structural categories [20, 21]. It also simulates orthogonal data supporting both known and novel transcripts.

SQANTI-SIM requires as input data the reference genome and the GTF file with transcriptome annotation for the organism under analysis. Users can define various parameters to control the novelty component of the simulated reads, such as the number, SQANTI3 structural category, and expression levels of the transcripts they wish to simulate. Optionally, long-read, short-read, and CAGE peak datasets can be provided to estimate read properties and simulate supporting orthogonal data. SQANTI-SIM returns the simulated long-reads, a reduced GTF file without the simulated novel transcripts, and the orthogonal datasets. Moreover, it includes functions to generate a comprehensive report that evaluates the performance of the transcript reconstruction algorithm when applied to the simulated data.

The workflow of SQANTI-SIM usage (Fig. 1) consists of the following steps:

1. Transcripts annotated in the reference GTF are classified according to their potential SQANTI3 structural category when compared to other transcripts of the same gene.
2. Based on this classification, a user-defined number of transcripts classified as ISM, NIC, NNC, or other novel structural categories, are removed from the annotation resulting in a “reduced” GTF file. Additionally, SQANTI-SIM assigns transcript expression values by offering three computation modes. The *equal* mode assigns the same expression value to all simulated transcripts. The *custom* mode allows the user to define different expression values for novel and known transcripts by customizing the parameters of two negative binomial distributions. Lastly, the *sample* mode utilizes inverse transform sampling from an empirical raw counts distribution.
3. Long reads are simulated using either NanoSim, PBSIM3, or IsoSeqSim operating on the complete GTF annotation file. Moreover, SQANTI-SIM can optionally simulate matching short reads and CAGE-peak data taking parameters from suitable reference datasets.
4. The transcriptome reconstruction algorithm utilizes the simulated reads and the “reduced” reference annotation generated by SQANTI-SIM to predict transcript models. If the transcriptome reconstruction algorithm permits, the simulated orthogonal data may be also incorporated into the transcript model prediction.
5. The performance of the method is assessed for each novel structural category using the SQANTI-SIM evaluation function that identifies true and false novel transcripts based on the simulated ground truth. Transcripts missing in the reduced GTF should be identified as true novel, and any novel transcript that was not simulated will result in a false call. A full performance evaluation report is provided.

**Fig. 1.**
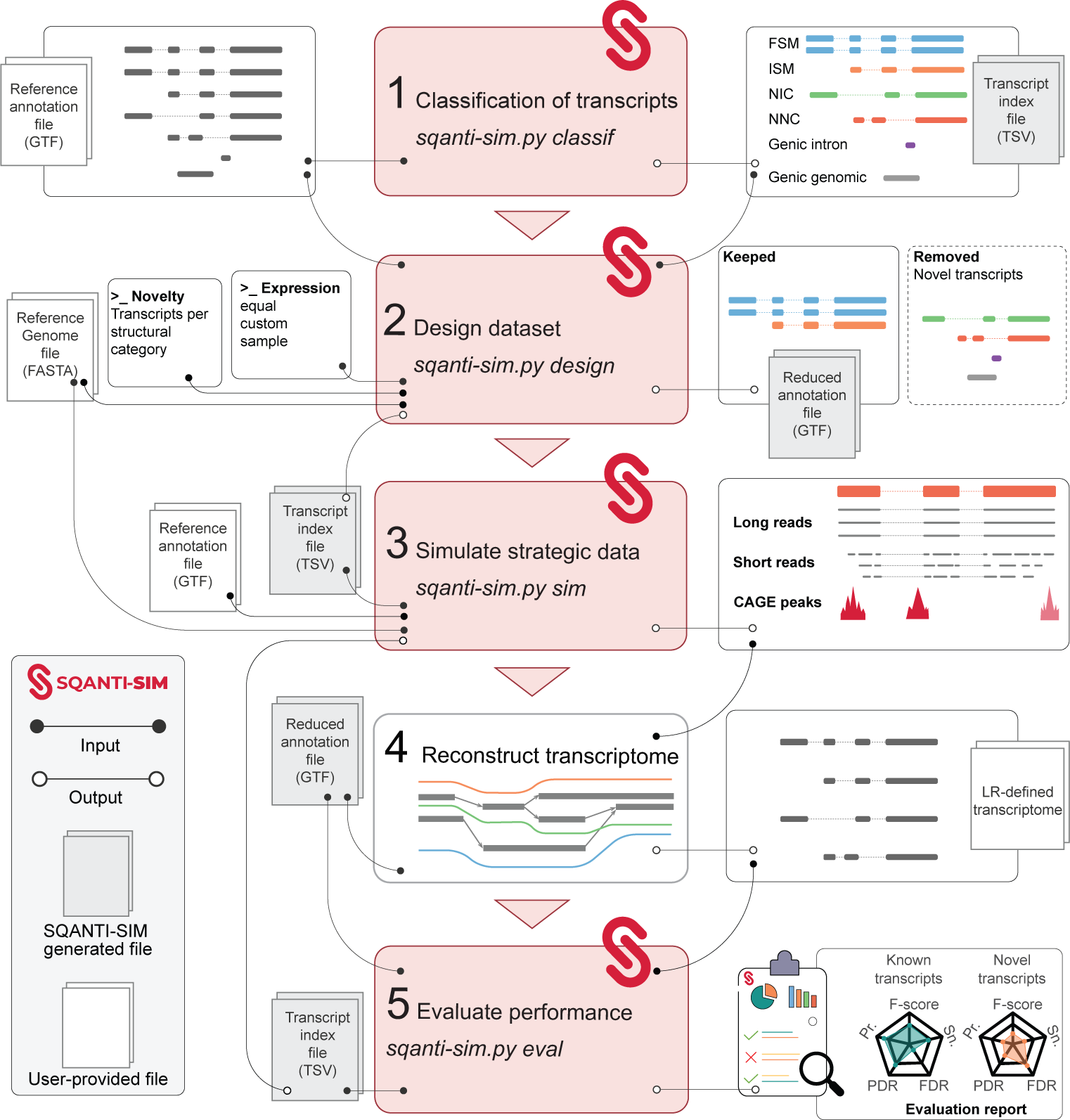
Flowchart of the SQANTI-SIM pipeline. The first three steps simulate reads and accompanying datasets according to the user’s specifications. Simulated data is then used by the transcriptome reconstruction algorithm to predict transcripts. The last SQANTI-SIM module assesses performance by comparison to the simulated ground truth and provides a comprehensive evaluation report.

### 2.2 Validation of the SQANTI-SIM simulation approach

We demonstrate the reliability of the SQANTI-SIM framework by simulating cDNA ONT and cDNA PacBio long-read datasets with the *sample* mode. These simulations were based on the human WTC11 cell line lrRNA-seq data from the LRGASP project [17]. Furthermore, as Illumina and CAGE-seq datasets are available for the same cell line, SQANTI-SIM was tasked with simulating the corresponding Illumina and CAGE-peak data (detailed in the Methods section). We conducted an assessment of SQANTI-SIM’s ability to accurately simulate the desired transcript types and to faithfully mirror the empirical distributions of the sample data.

Since a notable feature of SQANTI-SIM is its ability to simulate novel transcripts, our initial assessment focused on the tool’s accuracy in effectively recreating various types of novel transcripts. The transcript number of each simulated SQANTI3 structural category and their assigned reads are provided in Supplementary Table 1. The simulation of novel transcript categories involves the coordinated elimination and retention of specific transcript annotations to establish the novelty status, a task that might prove challenging for genes with many transcripts. For this validation, we requested 4300 FSM and 1000 transcripts of each novel SQANTI3 category to be simulated. To verify the correct simulation of novelty, we ran SQANTI3 analysis on the reduced GTF and confirmed that all structural categories were simulated in the requested amounts (Fig. 2a), while still preserving the number of transcripts per gene of the real dataset (Fig. 2b). This result reveals that SQANTI-SIM is able to control the transcript novelty during simulation while maintaining the complexity of the reference transcriptome.

**Fig. 2.**
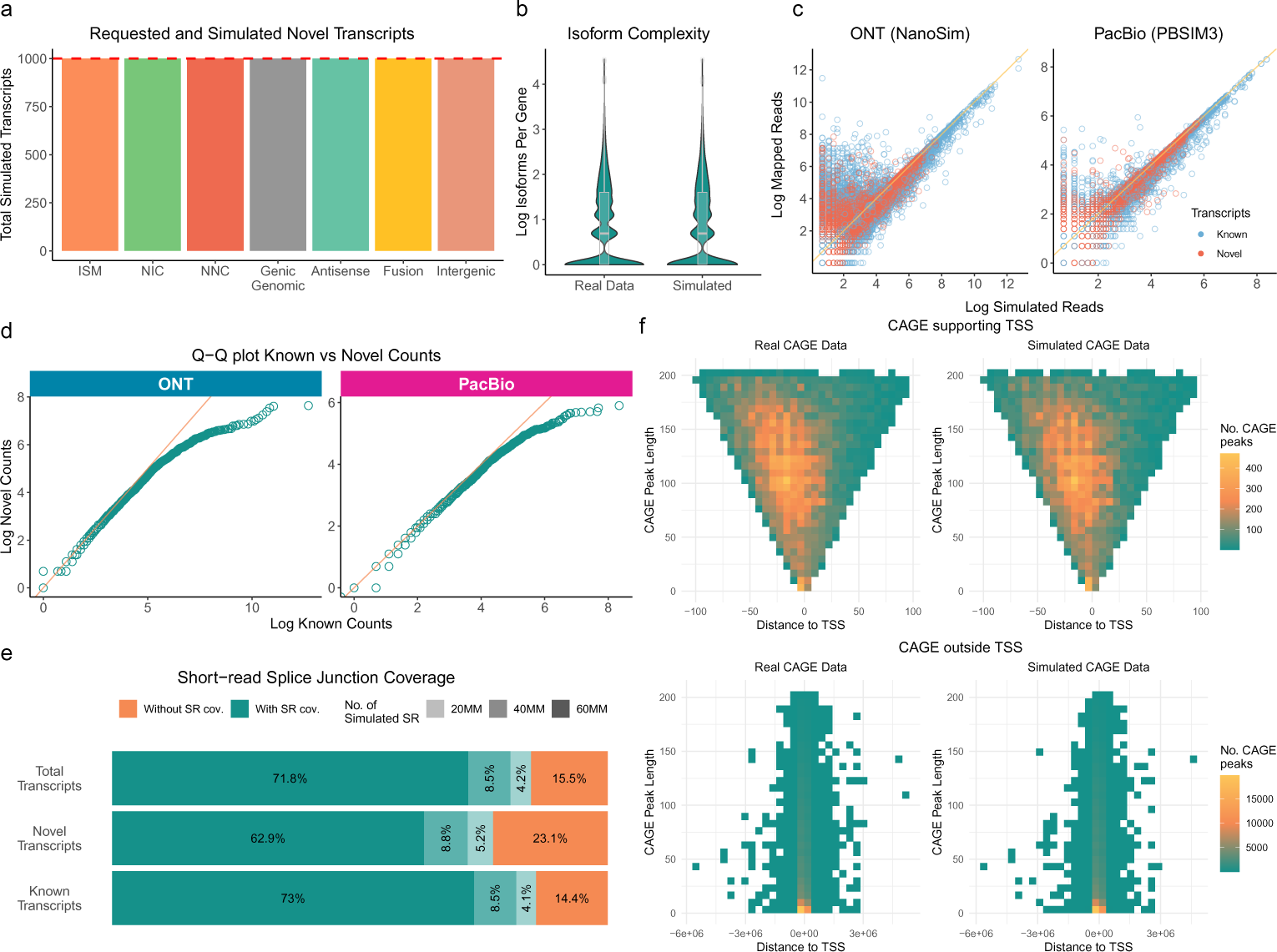
Validation of SQANTI-SIM approach. **a** Simulated transcripts for each novel SQANTI3 structural category out of 1000 requested for each type (red line). **b** Comparison of the distribution of expressed isoforms per gene in real human WTC11 cell line data and simulated data. **c** Scatter plot comparing simulated transcript reads and mapped raw counts. **d** Quantile-Quantile (Q-Q) plot comparing the expression levels, represented as log raw counts, between known and novel transcripts. **e** Proportion of simulated transcripts with all splice junctions (SJ) covered by sample-specific short-read (SR) data at different simulated sequencing depths. **f** Bivariate distribution of CAGE peak length and distance to the closest transcript transcription start site (TSS) for CAGE peaks supporting a TSS and those not supporting any TSS in real WTC11 and simulated data.

To assess the accuracy of expression value simulation, we employed minimap2 to map reads and then compared the number of primary alignments obtained with the number of simulated reads for each transcript. We found a high correlation between the simulated and the mapped data for both read simulators, although NanoSim produced more noisy reads (lower mappability) than PBSIM3, as expected given the differences in error models between the two sequencing platforms (Figure 2c). For this validation, the simulation of novel transcripts included the utilization of the *–diff exp* option available within the *sample* mode. This option permits users to define the bias between expression levels of novel and known transcripts. Larger values result in novel transcripts having lower expression. We confirmed that SQANTI-SIM successfully simulated lower expression values for novel transcripts (Figure 2c) and achieved the intended difference in overall expression value distributions between known and novel transcript models (Figure 2d).

In addition to long-read data simulation, SQANTI-SIM simulates matching orthogonal data in the form of short reads and CAGE peaks, providing a multi-omic simulation dataset where supporting data can effectively be used for transcript model inference. SQANTI-SIM utilizes Polyester for short-read data simulation and provides the algorithm with the same transcript expression level distributions as for the long reads to obtain a compatible dataset. We validated this strategy by confirming that short reads simulated at an increasing sequencing depth increasingly supported long-read transcript model splice junctions, with junctions in known transcripts being more frequently supported than those in novel transcripts, as expected due to their lower simulated expression values (Fig. 2e).

Finally, SQANTI-SIM simulates matching CAGE peaks from short reads by applying a logistic regression model. Users may use the SQANTI-SIM-included pre-trained model obtained with the WTC11 cell line data or provide their own sample-specific CAGE-peak dataset. Moreover, users can specify the desired proportion of CAGE peaks that overlap a transcription start site (TSS) to be simulated or obtain this value from their data. SQANTI-SIM employs inverse transform sampling to determine the length and genomic position of each CAGE peak, based on empirical bivariate distributions. Using the WTC11 dataset we confirmed that SQANTI-SIM CAGE-peak simulation faithfully recapitulates empirical data both for CAGE peaks that support and do not support transcript TSS (Fig. 2f).

### 2.3 Application to benchmark transcriptome reconstruction methods

In order to demonstrate the utility of SQANTI-SIM as a benchmarking tool, we simulated cDNA ONT and PacBio datasets containing FSM, ISM, NIC, and NNC transcripts (Supplementary Table 2) and used these data to analyze the performance of widely used transcriptome reconstruction algorithms. We selected two reference-based methods: FLAIR (with and without short-reads data usage) and TALON, and the PacBio-official reference-free IsoSeq approach alone and in combination with SQANTI3 Rules filter [20], making a total of five different transcript reconstruction pipelines (TALON, FLAIR, FLAIR+Illumina, IsoSeq, and IsoSeq+SQ3Rules) for evaluation. As IsoSeq is not compatible with Nanopore data, this method was only applied to the simulated cDNA PacBio dataset.

The predicted transcriptomes obtained with the different analysis pipelines exhibited a large variation in the number of detected transcripts, the ratio between True Positives (TP) and False Positives (FP), and the distribution of structural categories, despite being derived from the same datasets (Fig. 3a). TALON and FLAIR reference-guided approaches predominantly yielded known FSM transcript models, except for TALON processing ONT data that included an important fraction of ISM. In contrast, the reference-free IsoSeq pipeline identified over 60% novel transcripts, and this percentage was substantially reduced after applying the SQANTI3 Rules filter. These results reveal distinct strategies employed by lrRNA-seq reconstruction algorithms to manage the novelty of transcript models.

**Fig. 3.**
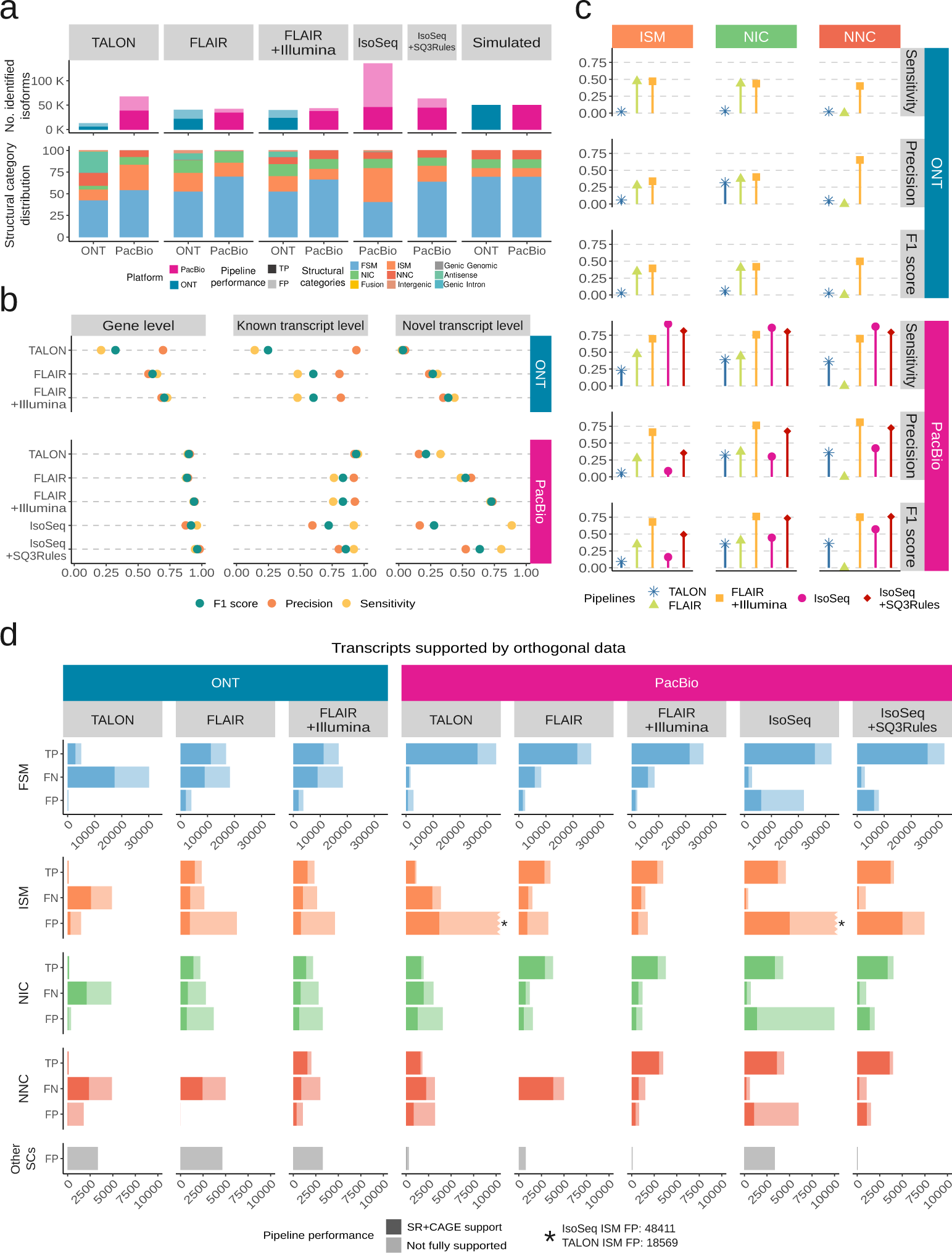
Benchmark of isoform identification pipelines. **a.** Distribution of SQANTI3 structural categories and the number of detected true (TP) and false positive (FP) transcripts. **b** Performance of analysis pipelines in the detection of known and novel simulated transcripts. **c** Performance based on different types of simulated novelty (ISM, NIC, and NNC). **d** TP, FN and FP transcripts for each pipeline and structural category, with and without support from both SJ short-read coverage (min cov ¿ 1) and TSS with CAGE peak support.

SQANTI-SIM provides evaluation metrics at both gene and isoform levels. While gene-level predictions demonstrated strong performance on the PacBio dataset, the accuracy of transcript model estimates was comparatively lower (Fig. 3b). At the transcript level, reference-guided pipelines exhibited diminished accuracy and sensitivity in identifying novel transcript models in comparison to those already annotated. The reference-free IsoSeq demonstrated high sensitivity and low precision both for known and novel transcripts, IsoSeq+SQ3Rules substantially increased precision and F1 score and FLAIR+Illumina, combining long and short reads, had the highest F1 scores for gene, known, and novel transcript predictions. Overall, the ONT predictions displayed poorer precision and sensitivity, both for known and novel transcripts in all tested methods (Fig. 3b).

Since SQANTI-SIM simulates the SQANTI3 transcript structural categories in a controlled manner, we can evaluate how different analysis pipelines handle the various types of transcript novelty (Fig. 3c and Supplementary Fig A1). We found that FLAIR solely using long reads had remarkably lower sensitivity and precision for NNC and a slightly worse precision for ISM compared to NIC transcripts, all of which were greatly recovered when short reads were added to the prediction. This suggests that FLAIR improves novel transcript estimates when using short reads data through an improved calling of novel splice-sites and transcript ends. TALON, on the contrary, using cDNA PacBio simulations, struggled more with the correct identification of ISM than with other novelty types. Finally, IsoSeq showed a high sensitivity for all the evaluated novel transcripts types, but particularly low precision for ISM, indicating a large number of false positives in this category. This low precision of IsoSeq was notably increased after applying the SQANTI3 Rules filter, especially for the NIC and NNC transcripts. Overall, SQANTI-SIM reveals that the ISM category exhibits the poorest performance across all analysis methods and sequencing platforms (Fig. 3c), highlighting the challenges that lrRNA-seq faces to accurately define TSS and/or TTS.

Given the availability of simulated orthogonal data from SQANTI-SIM, we are able to assess the degree to which these data support transcript model predictions for each structural category (Fig. 3d). Our observations indicate that, in general, transcript models predicted from ONT data exhibit less orthogonal support compared to those predicted using PacBio reads. As expected, TP transcripts received substantial support from Illumina and CAGE-peak data across all structural categories. Remarkably, FN transcripts also displayed consistently high orthogonal support. This suggests that the inability of prediction tools to identify these transcripts is not linked to a reduced likelihood of being corroborated by additional data. Conversely, FP transcripts were predominantly found among those lacking orthogonal support, particularly evident in TALON and IsoSeq predictions and specifically within the ISM category. This implies that incorporating additional data support into these methods would lead to a reduction in FP rates, a characteristic demonstrated effectively by the IsoSeq+SQ3Rules pipeline, which implements filtering based on external data. Nevertheless, it’s worth noting that numerous FP ISM calls did possess both CAGE-peak and Illumina support. This underscores once again the challenge presented by this type of transcript for accurate identification in human lrRNA-seq datasets.

Finally, our controlled simulation approach allowed us to investigate the factors driving differences in the detection of transcripts by the evaluated pipelines. We compared the expression level, transcript length, and number of exons of TP, FP, and, FN transcripts in each structural category detected by each pipeline. We observed that for many pipelines, FN transcripts were longer (Supplementary Fig. A2a) and had a higher exon count (Supplementary Fig. A2b) compared to those that were successfully identified, especially considering the NIC category, suggesting that accurately detecting lengthy transcripts could still pose challenges for certain lrRNA-seq methods. Interestingly, this pattern did not hold true for TALON when applied to ONT data. In this case, FN transcripts were notably shorter than true positive (TP) ones, revealing a distinct behavior of this tool on the Nanopore platform. However, a more consistent pattern emerged when examining expression levels (Supplementary Fig. A2c). TP transcripts consistently exhibited higher expression levels compared to FN transcripts, particularly in the case of Nanopore data. This finding underscores the significance of expression levels as a crucial factor when generating transcript models with all the examined analysis methods.

## 3 Discussion

The long-read RNA-seq technology has significantly enhanced our ability to profile transcriptome complexity and has facilitated the study of isoform diversity across a broad spectrum of organisms. This advancement has led to the unearthing of thousands of novel isoforms, even within extensively studied species [30–32]. However, benchmarking studies of lrRNA-seq have shown notable discrepancies in the identification of novel transcript models among different sequencing platforms and analysis methods [2, 7, 17–19]. This disparity underscores the intricate challenge of accurately identifying novel transcripts from long-read sequencing data. As the adoption of lrRNA-seq for transcriptome analysis continues to expand and the number of reported isoforms dramatically increases, it becomes imperative to have access to tools capable of facilitating the precise assessment of novel transcript calls.

SQANTI-SIM has been developed as an effective tool to offer a reliable framework for simulating novel transcripts in a controlled manner. By making use of the structural category classification of SQANTI3 and real transcript models from the reference annotation, SQANTI-SIM generates simulated transcriptome annotation files where both known and different types of novel isoforms are defined, and employs well-established long-read simulators (NanoSim, PBSIM3, and IsoSeqSim) to simulate reads. This approach offers several advantages. Firstly, the simulated novel transcripts correspond to authentic transcript structures, rendering them more credible compared to alternatives that generate synthetic or artificially constructed transcripts through exon merging or transcript alignment across diverse species. Artificial transcripts, aside from possibly introducing unrealistic novel events, might lack validation from orthogonal data, frequently employed to support transcript predictions. In line with this notion, SQANTI-SIM incorporates the simulation of short-read and CAGE-peak data tailored to the simulated long-read dataset. These datasets can serve a dual purpose: to evaluate the potential validation by orthogonal data for the newly predicted transcripts or to directly aid in transcript reconstruction methods that accept this form of supplementary information. As far as our knowledge extends, SQANTI-SIM represents the first multi-omics simulation strategy designed for lrRNA-seq studies.

Secondly, SQANTI-SIM operates at the level of the definition of transcript models rather than on the read simulation task, for which established algorithms are used. This ensures a unified framework for simulating novel transcripts for both Nanopore and PacBio data, offers users the flexibility to choose their preferred read simulation algorithm, and facilitates the integration of new long-read simulators as they become available. Finally, SQANTI-SIM grants precise control over the quantity and expression levels of diverse novel transcripts categorized according to the widely-accepted SQANTI3 structural categories. This empowers users to assess how different reconstruction methods perform in identifying transcripts with novel splice sites, alternate transcription termination sites (TTS) or transcription start sites (TSS), or those that are intergenic. Additionally, by regulating the expression level of each known or novel transcript type, users can explore the boundaries of novel transcript detection, particularly when these transcripts are expressed at low levels. Moreover, due to SQANTI-SIM’s ability to maintain transcriptome complexity while simulating the reduced reference, the evaluation of novel transcript detection can be conducted within the accurate context of transcriptome diversity.

The SQANTI-SIM framework may have some limitations. It relies on the existence of a reliable reference annotation where known and novel transcripts can be defined. While this is likely a common requirement for any simulation strategy, it restricts application to well-annotated species, and users working with poorly annotated organisms may require to use well-characterized related species to construct their simulated datasets. Furthermore, the accuracy of the long-read simulation is reflective of the underlying algorithms used for simulating sequencing errors and library construction artifacts. This accuracy can vary depending on the chosen simulation tool. Lastly, the simulation of orthogonal data is contingent upon the presence of corresponding datasets. While this is often the case for short reads, CAGE data might not be available for many species. Nevertheless, this does not preclude the utilization of SQANTI-SIM’s novelty simulation, which does not necessitate any complementary dataset. Alternatively, users can consider using the pre-computed models of human Illumina and CAGE peak provided by SQANTI-SIM for a different approach.

To showcase the utility of SQANTI-SIM, we employed the tool to simulate ONT and PacBio cDNA data. We then evaluated various long-read RNA-seq reconstruction algorithms, which vary in their utilization of reference annotation and reliance on additional information. Our findings reveal significant disparities among these methods in accurately defining novel transcripts, with superior performance observed with simulated PacBio reads over ONT reads. SQANTI-SIM highlighted the interplay of sensitivity and precision across different SQANTI3 structural categories and analysis tools. For instance, while IsoSeq exhibited high sensitivity in detecting all transcript types, its precision was notably lower for ISM compared to other novel transcripts. Similarly, TALON’s accuracy was poorer for ISM than for other categories. These results suggest extra challenges in distinguishing these novel isoforms from partial sequences. Moreover, the SQANTI-SIM framework illuminated the substantial advantages of integrating orthogonal data into transcript model definitions. For example, FLAIR struggled to identify NNC transcripts without short-read support, but accuracy significantly improved with the inclusion of Illumina reads, as these may help in the precise delimitation of novel transcript and exon boundaries. Similarly, by enforcing CAGE and Illumina support through SQANTI3 Rules filtering, IsoSeq reduced false positives for ISM and NIC, resulting in a remarkable enhancement in performance for these transcript types. Furthermore, SQANTI-SIM’s ability to precisely control transcript novelty, structure, and expression level facilitates investigating potential factors contributing to inaccurate transcript identification. Our analysis revealed that transcript expression level constitutes a primary source of both false negatives and false positive, revealing present limitations of lrRNA-seq technologies for accurately defining transcripts expressed at low levels.

In conclusion, the discovery of novel transcripts through LRS remains a challenging task. We demonstrate that SQANTI-SIM stands as an essential and cost-effective resource for benchmarking lrRNA-seq technologies and analysis tools, addressing an important gap in the long-read sequencing field.

## 4 Methods

### 4.1 SQANTI-SIM pipeline

The SQANTI-SIM pipeline is implemented in Python and makes use of R [33] for specific functionalities such as short-read simulation and evaluating reconstructed transcriptomes. SQANTI-SIM is a wrapper script (*sqanti-sim.py*) consisting of distinct modules that perform tasks in the following order: (i) the *classif* stage screens the provided reference GTF, classifying annotated transcripts based on their potential SQANTI3 structural category compared to other transcripts of the same gene; (ii) the *design* generates a reduced GTF annotation file, excluding novel isoforms according to user-specified novelty types and amounts; (iii) the *sim* step runs NanoSim, PBSIM3 or IsoSeqSim to simulate reads based on the complete reference annotation, using the expression value distributions provided by the user; and the (iv) *eval* module assesses the accuracy of the long-read reconstructed transcriptome by running SQANTI3 for the predicted transcripts using the reduced GTF and an additional transcript index file containing the structural annotation of the simulated novel transcripts. For convenience, the *classif*, *design* and *sim* modules can be run in a single command called *full-sim*, facilitating full dataset simulation. We also provide pre-trained models and characterized datasets generated using the WTC11 human cell line that can be directly used without the need for additional training or user-provided files.

All results presented in this manuscript were generated using version 0.2 of SQANTI-SIM. For a more detailed explanation of SQANTI-SIM, please refer to our GitHub repository: https://github.com/ConesaLab/SQANTI-SIM.

#### 4.1.1 SQANTI-SIM classification

The *classif* module categorizes reference transcripts using the same classification algorithm as in the latest SQANTI3 version (v5.1.2. at the time of writing this work), providing consistent transcript structural classification across SQANTI3-based tools. Each transcript is assigned a structural category when compared to other transcripts of the same gene. We exclude self-comparisons which would always result in FSM.

SQANTI-SIM *classif* generates an index file containing reference transcript identifiers, their potential structural categories with the corresponding associated transcript, the associated gene, and other structural features such as length and number of exons. The classification of transcripts into potential SQANTI3 categories is the basis for the selection of novel transcripts for simulation according to user preferences. The resulting file is utilized as input for the subsequent SQANTI-SIM modules.

#### 4.1.2 SQANTI-SIM novelty selection

SQANTI-SIM *design* performs two main tasks: generating a reduced transcriptome reference annotation and setting expression levels. Reference transcripts are classified as “novel” or “known” based on user-defined requests for each structural category. “Novel” transcripts are removed from the annotation, resulting in a reduced GTF file, while retaining the associated reference gene/transcript information to preserve the previously assigned structural category. By default, SQANTI-SIM *design* targets only transcripts with a minimum length of 200 bp to avoid simulating small RNAs.

After selecting transcripts for simulation, expression values are assigned. SQANTI-SIM provides three alternatives to setting expression values. In the *equal* mode, the same expression value is assigned to all simulated transcripts, regardless of their novelty type. The *custom* mode allows for the customization of separate negative binomial distributions for known and novel transcripts, to introduce differences in expression levels between the two transcript types. The *sample* mode obtains expression values from a real expression distribution profiled from an existing long-read RNA-seq sample. Minimap2 [34] is used to obtain primary alignments to the reference transcriptome that is generated from the genome fasta file and the reference GTF using the gffread tool [35]. Inverse transform sampling is applied to simulate values from the empirical distribution. Minimap2 is run with different presets for Pacific Biosciences (PacBio) reads (”*-x map-pb*”) and for Oxford Nanopore Technologies (ONT) reads (”*-x map-ont* “). To maintain the possibility for different expression levels between known and novel transcripts while using the *sample* mode, the *–diff exp* parameter is used in the random sampling process to control the bias between the expression distributions of novel and known transcripts. The *–diff exp* parameter influences the odds of assigning high expression values to these two types of transcripts, with higher values resulting in a stronger bias towards known over novel transcripts. Additionally, relying on the primary long-read alignments, SQANTI-SIM roughly estimates the number of transcripts expressed for each gene. The *–iso complex* parameter forces SQANTI-SIM to simulate the diversity of transcripts observed, reproducing the isoform complexity of the sample.

Pre-computed empirical cumulative distribution functions (ECDFs) of expression values are provided by default for cDNA and dRNA ONT samples and a cDNA PacBio sample from the WTC11 cell line.

#### 4.1.3 SQANTI-SIM simulation

The SQANTI-SIM *sim* tool simulates reads from transcripts that were selected at the *design* stage. For long-read simulation, NanoSim, PBSIM3, or IsoSeqSim are used as detailed in Supplementary Methods Section 1. Users can either provide their own training data or used SQANTI-SIM pre-trained models created using the human WTC11 cell line data from the LRGASP project.

NanoSim v3.1.0 is employed for simulating cDNA and dRNA ONT reads. The execution of NanoSim includes the options “*-b guppy –no model ir* “ and controls the number of simulated reads by disabling the simulation of randomly unaligned reads. SQANTI-SIM simulates cDNA PacBio reads using either PBSIM3 v3.0.0 (*–pbsim*) or IsoSeqSim v0.2 (*–isoseqsim*). PBSIM3 is set to simulate the generation of CLR by multi-pass sequencing using the error models constructed from the provided PacBio reads, and *ccs* [36] is executed with default parameters to generate HiFi reads. For IsoSeqSim, normal mode is employed, utilizing the 5’ end and 3’ end completeness information from the provided pre-trained PacBio Sequel model, along with the user-defined error rates.

Additionally, SQANTI-SIM simulates matching complementary data such as short-read Illumina data and CAGE-peak data. Illumina RNA-seq reads are simulated using the Polyester R package [37] and expression values are assigned using the same TPMs as for long-read data, resulting in two FASTA files with paired-end reads of 100 nt in length. These simulated reads are generated using an error model uniformly distributed at an error rate of 0.5%.

When used in *sample* mode, SQANTI-SIM provides a data-driven approach for simulating CAGE-peak orthogonal data based on short-read expression (Supplementary Fig. A3). SQANTI-SIM starts by estimating CAGE-peak length and determining the distance from each CAGE-peak center to the nearest transcript TSS using the user-provided CAGE-peal BED file. This results in two separate empirical bivariate distributions (one for length and another for distance) for CAGE peaks that either overlap or do not overlap a transcript TSS.

To predict the presence or absence of a CAGE-peak supporting a given transcript TSS, SQANTI-SIM fits a logistic regression model. The first step involves computing the TSS ratio SQANTI3 metric and the proportion of TSS coverage using BEDTools (Supplementary Fig A3). The TSS ratio is the ratio of short-read coverage 100bp downstream and upstream of the TSS defined in the SQANTI3 software [20]. The proportion of TSS coverage is computed by considering a 20bp window downstream of the TSS and dividing by the total number of reads. These values, along with information on the presence or absence of a CAGE peak supporting each transcript TSS (*Y_i_* for each transcript *i* = 1*, …, n*, where *n* represents the total number of transcripts), are used to build the logistic regression model as follows:

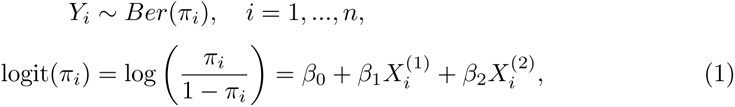

where *π_i_*is the probability of having a CAGE peak overlapping the transcript TSS*i* position, and *π_i_* is linked to the linear predictor by the *logit* link function. The linear predictor consists of an intercept term *β*_0_ and coefficients *β*_1_ and *β*_2_, which correspond to the log-transformed TSS ratio 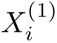 and the log-transformed proportion of TSS short-read coverage 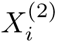, respectively.

The user can specify the proportion of CAGE peaks that should not align with any TSS or take this from the empirical data. Using the fitted logistic regression model and random sampling from the empirical cumulative distribution functions (ECDFs) of the bivariate distribution, SQANTI-SIM generates a BED file with the simulated CAGE peaks. SQANTI-SIM includes pre-trained models for CAGE-peak simulation obtained with the LRAGSP human WTC11 cell line data. This model achieved 81.2% accuracy in a 10-fold cross-validation

#### 4.1.4 SQANTI-SIM performance evaluation

The SQANTI-SIM *eval* module is used to assess the accuracy of the transcriptome reconstruction pipeline employed for identifying simulated transcripts. The evaluated transcriptome reconstruction algorithm should predict transcript models using the simulated data and the reduced reference annotation generated by SQANTI-SIM.

The *eval* step runs SQANTI3 to determine the structural categories of the reconstructed transcripts, utilizing the reduced annotation. Subsequently, SQANTI-SIM evaluates the accuracy of the reconstruction methods by comparing splice junctions and 3’/5’ transcript ends of the reconstructed transcript models with those present in the simulated transcripts. The potential structural categories and structural features of these simulated transcripts are stored in the SQANTI-SIM index file. SQANTI-SIM reports various statistics related to the accuracy of the retrieved data. Transcripts that were removed from the reference annotation should be identified as novel, and any new transcript not simulated by this controlled procedure is considered a false call. Sensitivity (Sn) and precision (Pr) are calculated at the isoform level for both novel and known transcripts and for each structural category separately:

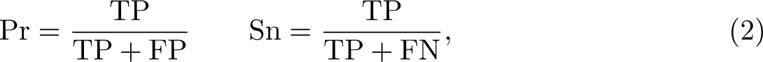

where true positive (TP) represents the reconstructed transcript models that match the simulated reference transcript at all splice junctions and have 3’/5’ ends within 50 nt from the annotated TSS and transcription termination site (TTS), false positive (FP) are transcripts that were detected but not simulated, and false negative (FN) indicates transcripts that were simulated but not identified according to the criteria for TP.

SQANTI-SIM also provides F_1_ scores as a summary measure of accuracy:

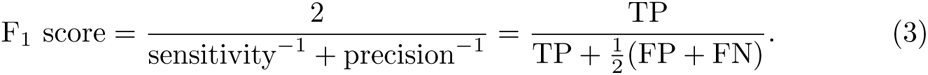

The evaluation report also includes metrics to evaluate partial matches. Partial true positives (PTP) are transcript models that match a reference transcript at all splice junctions but have a 3’/5’ end located more than 50 nts away from the TSS and/or TTS. Consequently, SQANTI-SIM calculates the Positive Detection Rate (positive detection rate (PDR)) and False Detection Rate (false discovery rate (FDR)) as follows:

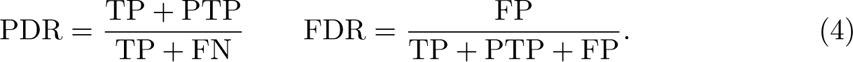

For comprehensive characterization of the TP, FN and FP calls, the SQANTI-SIM report included additional plots describing other structural features. These include transcript length, number of exons, number of simulated reads, and the number of transcripts with canonical or non-canonical junctions. Furthermore, provided with simulated orthogonal data, SQANTI-SIM *eval* generates descriptive plots, including visualizations of SJ coverage for TP and FP transcripts using short-read data, as well as the TSS support by CAGE peaks.

For an example, of the SQANTI-SIM evaluation report, we refer to https://github.com/ConesaLab/SQANTI-SIM/blob/main/example/example_SQANTI-SIM_report.html.

### 4.2 Real data characterization and dataset simulation

#### 4.2.1 Pre-trained models and empirical distributions for WTC11 in SQANTI-SIM

SQANTI-SIM, includes the default pre-trained models and empirical distributions utilized by the long-read simulators (NanoSim, PBSIM3, and IsoSeqSim), although users have also the option to define custom distributions. For the long-read simulator models, PBSIM3’s default choice is a quality score model derived from PacBio RS II reads provided by PBSIM3 developers. By default, IsoSeqSim is used with the error rates recommended in the IsoSeqSim GitHub repository, which are based on PacBio Sequel data (substitution 1.731%, deletion 1.090%, and insertion 2.204%). For NanoSim we provide specific profiles and pre-trained models for cDNA-ONT (ENCODE accession ENCFF338WQL) and dRNA-ONT (ENCODE accession ENCFF155CFF) reads acquired from the human WTC11 cell line sequencing data generated by the LRGASP project [17] (see Supplementary Methods Section 2.1). The average error rates for these pre-trained models are 8.1% for cDNA (mismatch 2.8%, insertion 1.9% and deletion 3.5%) and 12.2% for dRNA (mismatch 3.6%, insertion 3% and deletion 5.7%).

For expression value assignment at the *design* stage, SQANTI-SIM offers default empirical distributions of raw counts for various read types. Raw data associated with these distributions can be accessed through the ENCODE database, including cDNA-PacBio (accession ENCFF338WQL), cDNA-ONT (accession ENCFF263YFG), and dRNA-ONT (accession ENCFF155CFF). To obtain these empirical expression distributions, reads were mapped to the reference transcriptome using *minimap2* v2.26 (see Supplementary Methods Section 2.2), and primary alignments were computed as raw counts.

Additionally, SQANTI-SIM provides pre-trained models for simulating CAGE peaks. To create the models and profiles for CAGE-peak data, SQANTI-SIM requires long-read-defined transcript models and sample-specific short-read and CAGE peak data. SQANTI-SIM uses data from the WTC11 cell line available at the ENCODE database, with cDNA-ONT reads under experiment accession ENCSR539ZXJ, PacBio under accession ENCSR507JOF, short reads under ENCSR673UKZ, and CAGE peaks from GEO accession GSE185917. Transcript models were generated using IsoSeq v4.0.0 and FLAIR v2.0.0 [10], with assistance from matched short-read data (see Supplementary Methods Section 2.3.1). Short reads were aligned using STAR v2.7.10b [38] to obtain TSS coverage and the TSS ratio of reconstructed transcript models. Downloaded CAGE-peak calls from GEO were filtered to include only peaks found in at least 2 replicates, resulting in 139,285 CAGE peaks. CAGE peak data and model fitting were characterized using the supplementary script *sqanti-sim.py* in *train* mode (see Supplementary Methods Section 2.3.2).

#### 4.2.2 PacBio and ONT datasets simulation

##### SQANTI-SIM validation datasets

For validating the SQANTI-SIM pipeline we simulated cDNA ONT and PacBio reads including transcript models for all the SQANTI3 structural categories. A total of 50,000 isoforms were simulated, with 1,000 assigned to each novel structural category (Supplementary Table 1). The simulation was executed using SQANTI-SIM in *full-sim sample* mode with default *–long count* parameter. We used the *–iso complex* parameter to validate the simulation of the number of simulated transcripts per gene (see Supplementary Methods Section 3). Short-read data was simulated with various sample sizes (20M, 40M, and 60M short reads) using Polyester. The 20M short-read dataset was used to simulate CAGE-peak data.

##### Data simulation for pipeline benchmarking

The utility of SQANTI-SIM as a benchmarking tool was demonstrated by assessing the performance of different transcriptome reconstruction algorithms and pipelines. For this assessment, we simulated a cDNA ONT and PacBio dataset consisting of known isoforms (FSM) and novel transcripts falling into the three main SQANTI3 structural categories (ISM, NIC, and NNC) (see Supplementary Methods Section 4.1). Datasets were simulated using the GENCODE *H. sapiens* (GRCh38.p13) reference genome and GENCODE annotation v43. For both the PacBio and ONT datasets, SQANTI-SIM was executed in *–full-sim sample* mode, generating 50,000 different isoforms, out of which 15,000 were novel (5,000 ISM, 5,000 NIC, and 5,000 NNC). To simulate the ONT dataset, we used the parameters “*–ont –illumina –long count 20000000 –short count 40000000 –read type cDNA*”, which resulted in the simulation of 20 million ONT long reads and 40 million Illumina reads. For the PacBio dataset, we used the parameters “*–pb –pbsim –illumina –long count 4000000 –short count 40000000* “ to simulate 4 million PacBio reads with PBSIM3 and 40 million Illumina reads. We set the option “*–iso complex* “ to approximate the isoform complexity of the profiled real data, and “*–diff exp 2* “ to simulate a bias of lower expression for novel transcripts compared to known transcripts. We used the default provided pre-trained models and ECDFs from the WTC11 cell line in the simulation.

### 4.3 Long-read-defined transcriptome reconstruction methods

The performance of IsoSeq, IsoSeq+SQ3Rules filter, FLAIR, FLAIR+Illumina, and TALON was evaluated with the simulated datasets described in the previous section.

As IsoSeq [12] is designed to identify isoforms from reads sequenced on the PacBio sequencing platform, only the PacBio simulated data was processed with IsoSeq. The simulated HiFi reads were clustered using the *isoseq cluster2* mode. Then, reads were mapped to the reference genome using *pbmm2* v1.12.0 with the *ISOSEQ* preset. Finally, the alignments were collapsed using *isoseq collapse* with *–do-not-collapse-extra-5exons* option.

For the IsoSeq+SQ3Rules pipeline, the SQANTI3 v5.1.2 Rules filter was applied to the IsoSeq transcript models with the following parameters: Isoforms flagged for intrapriming or RT-Switching were discarded. FSM and ISM isoforms were required to have TSS support either from CAGE-peak overlap, having a TSS ratio of 1.5, or close proximity to annotated TSS (¡ 50bp). Additionally, other structural categories also required all splice-junctions to be supported by at least 2 short-reads or to be canonical junctions.

FLAIR [10] was applied to process both the ONT and PacBio datasets. Since FLAIR supports short-read splice data, it was run in two modes: (1) using only long reads and the “reduced” GTF (FLAIR pipeline), and (2) using long reads, the “reduced” GTF, and short-read splice sites for correction (FLAIR+Illumina pipeline). Short reads were mapped using STAR, and splice junctions with fewer than 3 supporting short reads were filtered from the resulting SJ.out.tab file. FLAIR was executed in the *123* mode, which performs alignment, correction, and collapse in a single step, using the “*–check splice*” parameter. Additionally, splice junction data was provided using the “*–short reads*” parameter when incorporating short-read simulated data.

Finally, TALON [11] was run on the ONT and PacBio mapped reads, guided by the “reduced” GTF. For the TALON pipeline, a database with the “reduced” GTF was generated using “*talon initialize database*” with the default settings. Then, the reads were mapped to the reference genome using minimap2 (*-ax splice:hq -uf –MD*), and “*talon label reads*” was used with the default settings to flag reads for internal priming. Next, the “*talon*” script was run to annotate the reads and novel transcript models were filtered using “*talon filter transcripts*”. Finally, the reconstructed transcriptome was built using “*talon create GTF* “.

Code details for all pipelines are provided at the Supplementary Methods Section 4.2.

**Supplementary information. Additional file 1:** Supplementary methods and code.

## Availability of data and materials

SQANTI-SIM is available at https://github.com/ConesaLab/SQANTI-SIM under GNU GPLv3 license. The human reference genome sequence (primary assembly, GRCh38) and gene annotation v43 from GENCODE can be obtained from https://ftp.ebi.ac.uk/pub/databases/gencode/Gencode_human/release_43. The long-read sequencing data obtained from the human WTC11 cell line is available at ENCODE (https://www.encodeproject.org/search): the cDNA ONT FASTQ file can be found under accession ENCFF263YFG, the dRNA ONT sample under file accession ENCFF155CFF, and the cDNA PacBio data under experiment accession ENCSR507JOF. Short-read sequencing data can be retrieved from accession number ENCSR673UKZ. The CAGE-Seq data is available at NCBI Gene Expression Omnibus (https://www.ncbi.nlm.nih.gov/geo/) under the accession number GSE185917. The simulated datasets generated and analysed during the current study can be reproduced using the code provided in the SQANTI-SIM v0.2 software repository and supplementary materials. Additionally, the files used to generate the results in this paper are publicly accessible at http://conesalab.org/SQANTI-SIM/.

## Authors’ contributions

A.C., F.P.P, and J.M.T. conceptualized and designed the SQANTI-SIM approach and the work. J.M.T. developed and implemented SQANTI-SIM, performed the analysis, and generated visualizations. T.L. contributed to implementing existing simulation tools in the SQANTI-SIM workflow. A.C. envisioned the study and supervised the work. J.M.T and A.C. drafted the manuscript. All authors read and approved the final manuscript.

## Tables

**Table 1.**
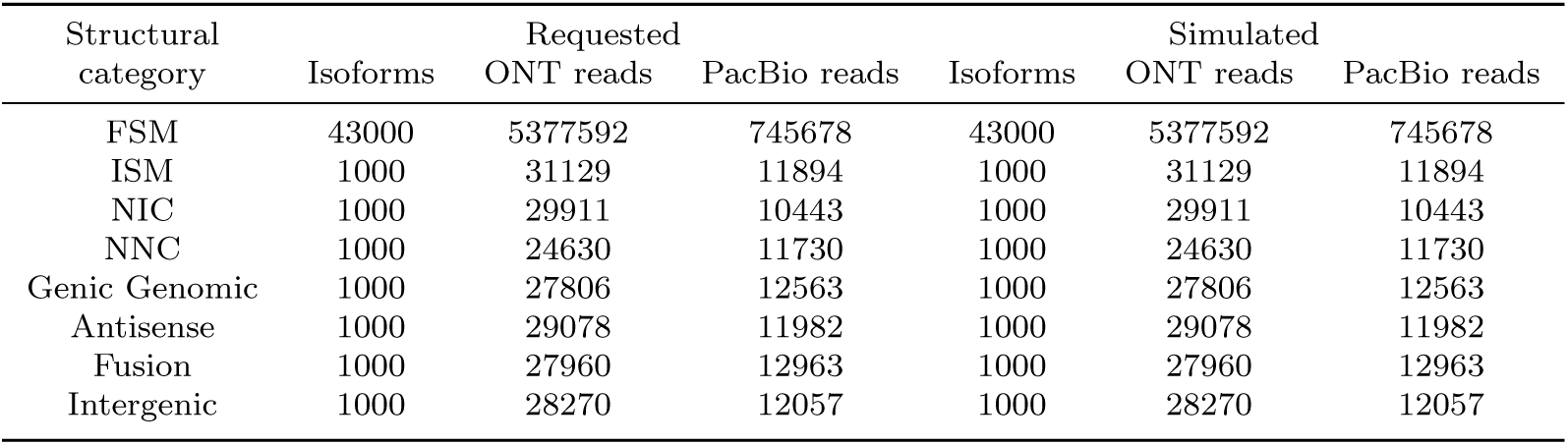
Requested and simulated transcript models and read counts for ONT and PacBio datasets used in SQANTI-SIM validation.

**Table 2.**
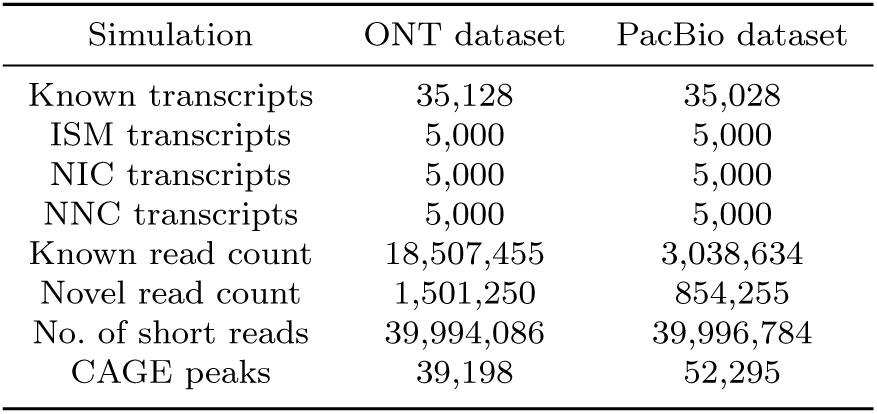
Simulated cDNA ONT and PacBio datasets used for pipeline benchmarking.

| Simulation | ONT dataset | PacBio dataset |
| --- | --- | --- |
| Known transcripts | 35,128 | 35,028 |
| ISM transcripts | 5,000 | 5,000 |
| NIC transcripts | 5,000 | 5,000 |
| NNC transcripts | 5,000 | 5,000 |
| Known read count | 18,507,455 | 3,038,634 |
| Novel read count | 1,501,250 | 854,255 |
| No. of short reads | 39,994,086 | 39,996,784 |
| CAGE peaks | 39,198 | 52,295 |

## Appendix A Supplementary Figures

**Fig. A1.**
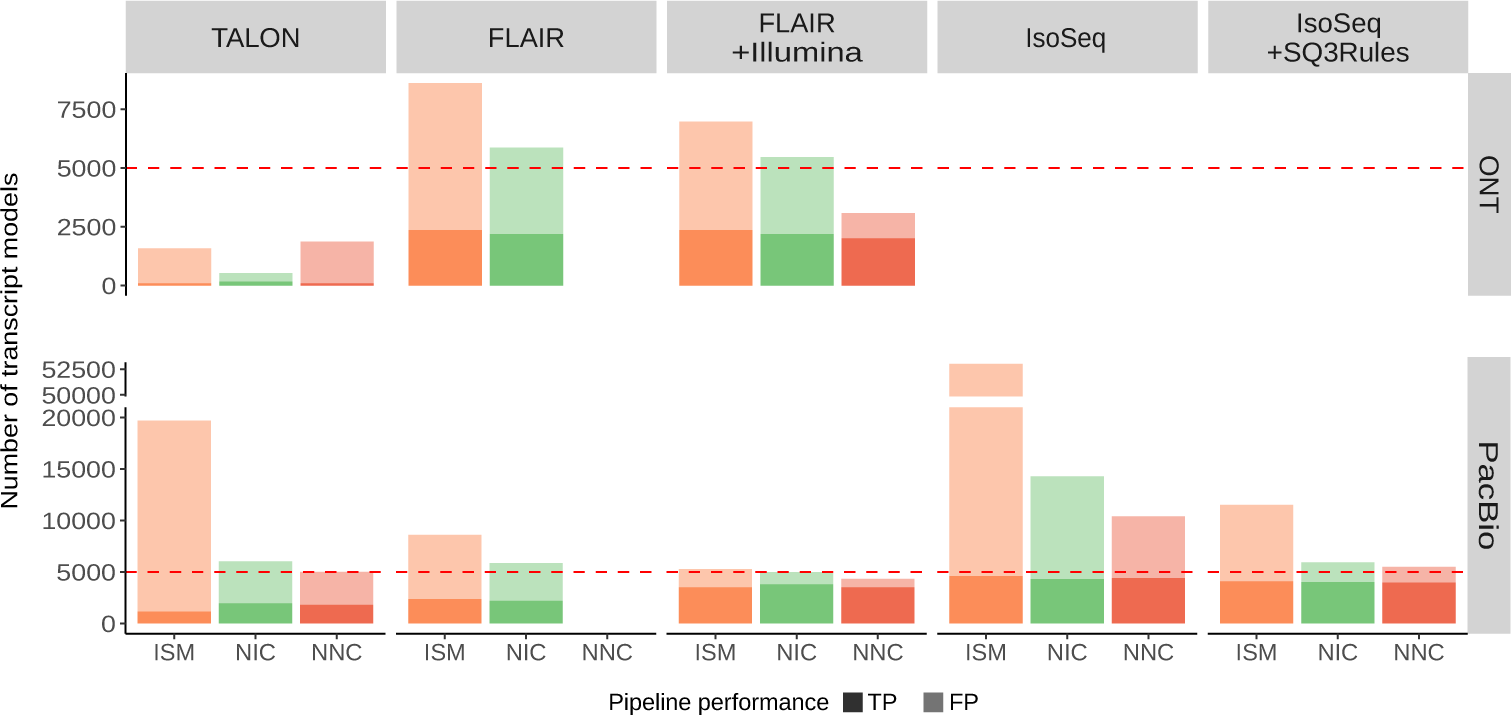
Number of detected true (TP) and false positives (FP) for different types of novelty (ISM, NIC, and NNC).

**Fig. A2.**
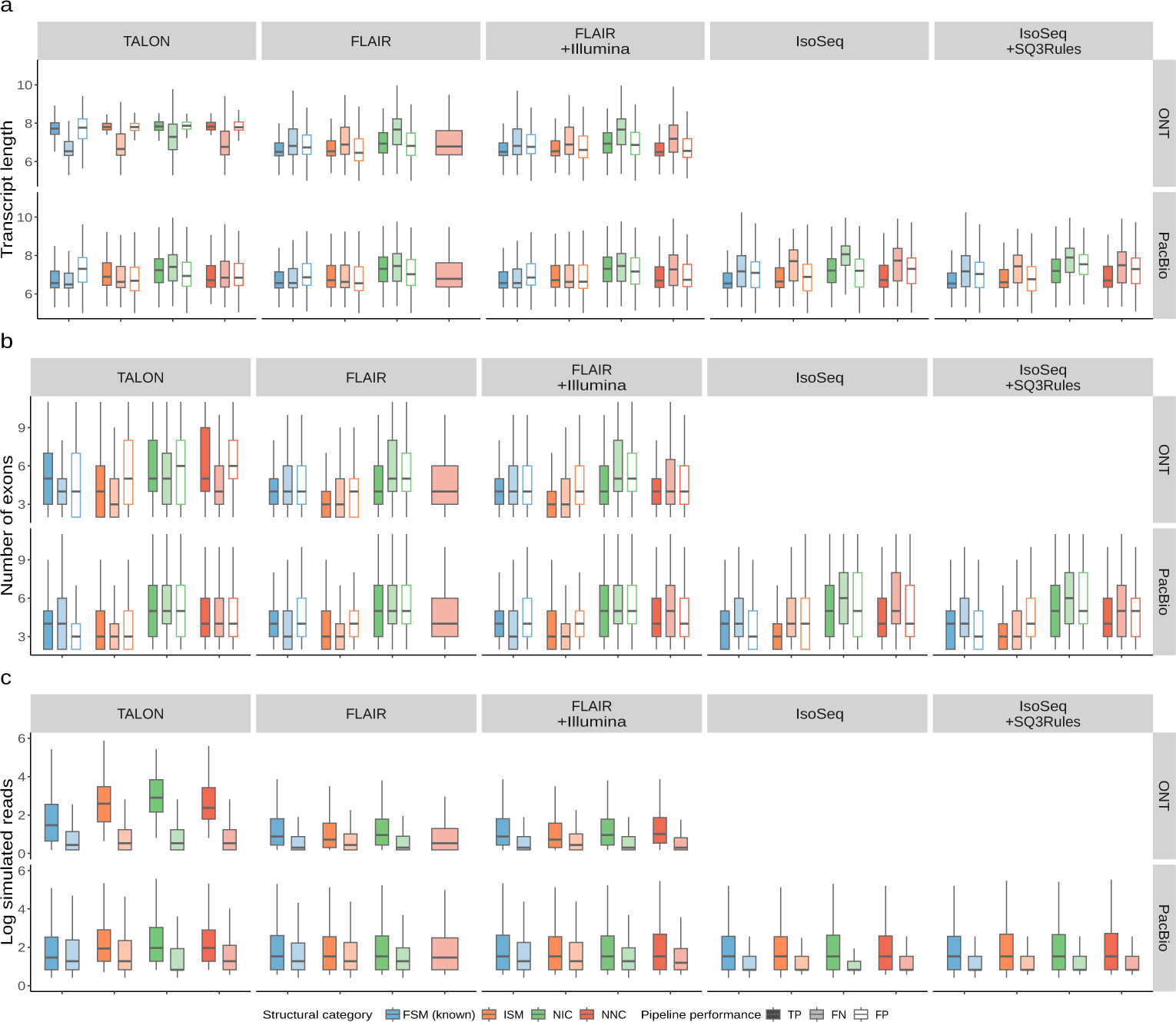
Relationship between true positives (TP), false negatives (FN), and false positives (FP) with (**a**) transcript length, (**b**) number of exons, and (**c**) simulated expression level.

**Fig. A3.**
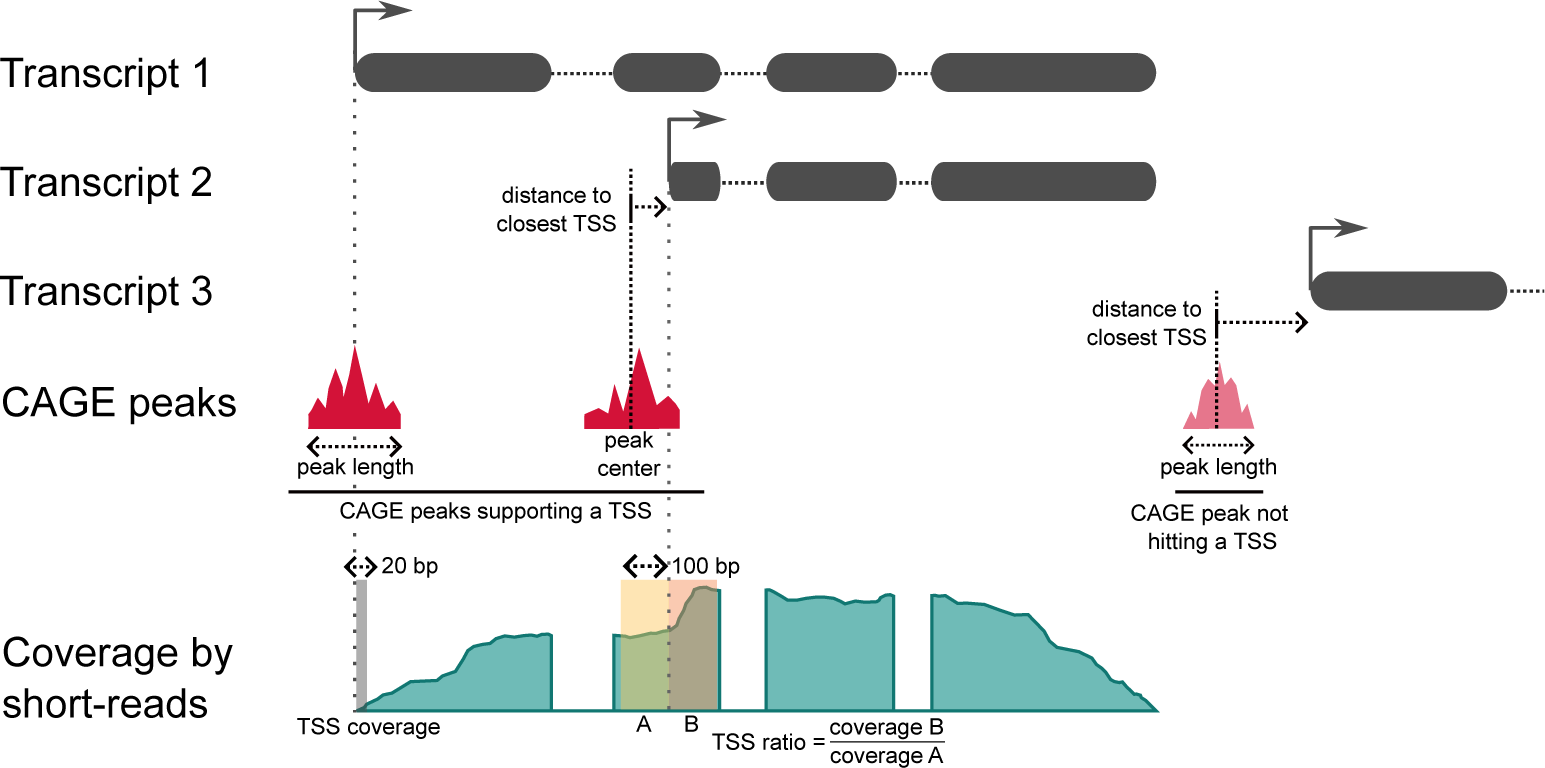
SQANTI-SIM characterization of CAGE peak data.

