## Supplementary methods for "SQANTI-SIM: a simulator of controlled transcript novelty for lrRNA-seq benchmark"

#### 1 SQANTI-SIM features

##### 1.1 PacBio simulation

When PacBio sequencing data is requested, SQANTI-SIM uses PBSIM3 as the default tool to simulate multi-pass transcriptome sequencing. This simulation process uses a quality score model. The following command is used to run PBSIM3:

```
pbsim \
    --strategy trans \
    --method qshmm \
    --qshmm data/QSHMM-RSII.model \
    --transcript <expression_profile> \
    --accuracy-mean 0.95 \
    --pass-num <number_of_passes> \
    --seed <seed>
```

If `--pass_num` $> 1$, the SQANTI-SIM pipeline executes CCS to generate PacBio HiFi reads:

```
ccs sd.bam sd.ccs.bam
```

Additionally, we have integrated IsoSeqSim's simulation within the SQANTI-SIM pipeline. If the `--isoseqsim` option is specified, IsoSeqSim will simulate PacBio Sequel data using the following command:

```
IsoSeqSim/bin/isoseqsim \
    -g <reference_genome> \
    -a <reference_annotation> \
    --expr <expression_profile> \
    --c5 utilities/5_end_completeness.PacBio-Sequel.tab \
    --c3 utilities/3_end_completeness.PacBio-Sequel.tab \
    -o <out_path/IsoSeqSim_simulated> \
    -t  <out_path/IsoSeqSim_simulated.tsv> \
    --es 0.01731
    --ed 0.01090
    --ei 0.02204
    -n <number_of_reads> \
    -m normal
    --cpu <num_cpus> \
    --tempdir <out_path/temp_isoseqsim> \
    --seed <seed>
```

##### 1.2 ONT simulation

If ONT sequencing data (cDNA/dRNA) is requested for simulation by SQANTI-SIM, it internally employs NanoSim to generate transcriptome reads through simulation. This process is carried out using the following command:

```
NanoSim/src/simulator.py transcriptome \
    -rt <reference_transcriptome> \
    -rg <reference_genome> \
```

```
    -e <expression_profile> \
    -c <error_profiles_from_characterization> \
    -o <out_path/ONT_simulated> \
    -n <number_of_reads> \
    -r {dRNA, cDNA} \
    -b guppy \
    -t <num_cpus> \
    --seed <seed> \
    --fastq --no_model_ir [--uracil]
```

##### 1.3 Illumina simulation

When short-read orthogonal data is requested for simulation, SQANTI-SIM internally utilizes Polyester to simulate a short-read RNA-seq paired-end sample. This simulation is performed using the following command:

```
simulate_experiment_countmat(
    <reference_transcriptome>,
    readmat = <expression_profile>,
    outdir = <out_path>,
    paired = TRUE,
    readlen = 100,
    seed = <seed>
)
```

#### 2 SQANTI-SIM WTC11 characterization

##### 2.1 NanoSim read characterization

We have updated NanoSim's error models by training them with more up-to-date sequencing data obtained from the LRGASP project.

```
annotation="gencode.v43.annotation.gtf"
genome="GRCh38.primary_assembly.genome.fa"
transcriptome="gencode.v43.transcripts.fa"

# ONT directRNA transcriptome reads
python NanoSim/src/read_analysis.py transcriptome \
        --read ENCFF155CFF.fastq \
        --ref_g $genome \
        --ref_t $transcriptome \
        --annotation $annotation \
        --aligner minimap2 \
        --output human_WTC11_dRNA_guppy_NanoSim/training \
        -t 8

# ONT cDNA transcriptome reads
python read_analysis.py transcriptome \
        --read ENCFF263YFG.fastq \
        --ref_g $genome \
        --ref_t $transcriptome \
        --annotation $annotation \
        --aligner minimap2 \
        --output human_WTC11_cDNA_guppy_NanoSim/training \
        -t 8
```

#### 2.2 Profile transcript expression pattern

In order to obtain the empirical isoform-level expression distribution for PacBio and ONT sequencing data, we aligned human WTC11 cell line long-read sequencing data to the reference transcriptome and retrieved primary alignments as transcript raw counts. The samples were obtained from ENCODE, and have the following file accession numbers: ENCFF338WQL (cDNA PacBio), ENCFF263YFG (cDNA ONT), and ENCFF155CFF (dRNA). Mapping of the reads to the reference transcriptome was performed using *minimap2*:

```
minimap2 \
    <reference_transcriptome> \
    <long_reads> \
    -x {map-pb, map-ont} \
    -a --secondary=no
    -o <out_sam_file>
    -t <num_cpus>
```

#### 2.3 Characterize CAGE-seq data

##### 2.3.1 Prepare input data

SQANTI-SIM requires LR-defined transcript models, along with matching short-read and CAGE peak data, to effectively characterize CAGE peak data and fit the model. We used human WTC11 cell line sequencing data generated by the LRGASP project. Long-read and short-read data can be accessed through the ENCODE database: cDNA-ONT reads are available under accession number ENCSR539ZXJ, cDNA-PacBio under experiment accession ENCSR507JOF, and the short reads under ENCSR673UKZ. CAGE-Seq data can be accessed through the Gene Expression Omnibus database under GEO accession GSE185917.
Short reads were mapped using STAR v2.7.10b. The resulting BAM files are used to later obtain TSS coverage and the ratio of TSS for transcript models.

```
genome="GRCh38.primary_assembly.genome.fa"
annotation="gencode.v43.annotation.gtf"
r11="wtc11_rep1.read1.fastq.gz"
r12="wtc11_rep1.read2.fastq.gz"

STAR \
    --runMode genomeGenerate --runThreadN 4 \
    --genomeDir $genome_dir --genomeFastaFiles $genome

# Repeat for all three replicates
STAR \
    --runThreadN 8 --genomeDir $genome_dir \
    --readFilesIn $r11 $r12 --outFileNamePrefix $out1 \
    --alignSJoverhangMin 8  --alignSJDBoverhangMin 1 \
    --outFilterType BySJout --outSAMunmapped Within \
    --outFilterMultimapNmax 20 --outFilterMismatchNoverLmax 0.04 \
    --outFilterMismatchNmax 999 --alignIntronMin 20 \
    --alignIntronMax 1000000 \ --alignMatesGapMax 1000000 \
    --sjdbScore 1 --genomeLoad NoSharedMemory \
    --outSAMtype BAM SortedByCoordinate --twopassMode Basic \
    --readFilesCommand zcat
```

For transcript model reconstruction, PacBio subreads were processed using IsoSeq v4.0.0, employing the *css* tool v6.4.0 with the ”*–min-rq 0.9*” parameter to generate ROIs. Primer removal and demultiplexing were performed using *lima* v2.7.1 with the options ”*–isoseq*” and ”*–peek-guess*”. Finally, FLNC reads were obtained after trimming the poly(A) tails and removing the concatemers with *isoseq refine* using the ”*–require-polya*” parameter.

```
ccs \
    $sampleName.bam \
        $sampleName.ccs.bam \
        --minLength 10 \
        --minPasses 3 \
        --min-rq 0.9 \
        --min-snr 2.5

primers="primers.fasta"

lima \
        $sampleName.ccs.bam \
        $primers \
        $sampleName.fl.bam \
        --isoseq \
        --peek-guess

 isoseq refine \
        $sampleName.fl.Clontech_5p--Clontech_3p.bam \
        $primers \
        $sampleName.flnc.bam \
        --require-polya
```

Transcript models for PacBio and ONT reads were identified using FLAIR, aided by matching short-read data. FLAIR v2.0.0 was employed with the "*–check_splice*" option, and the STAR SJ.out.tab file was provided to FLAIR using the "*–shortread*" parameter. We applied a filter to exclude splice junctions with less than 3 uniquely mapped reads to get high confidence splice junctions. Then, the filtered SJ.out.tab files were merged. FLAIR was then executed using the following command:

```
flair 123 \
    -r $cDNA_PacBio \
        -g $genome \
        -f $annotation \
        -o flair_LS_wtc11_pb \
        --temp_dir tmp_dir \
        --shortread wtc11_sr_SJ.filtered.merged.tab \
        --check_splice
```

##### 2.3.2 Profile CAGE data

To characterize CAGE peak data and fit the model, we used the FLAIR-defined transcriptome along with sample-specific short-read and CAGE-Seq filtered data from the WTC11 cell line.

```
isoforms="flair_LS_wtc11_pb.isoforms.chr.gtf"
genome="GRCh38.primary_assembly.genome.fa"
CAGE="ssCAGE/CAGE_WTC11.all_reps.merged.bed"
SR_bam="SR_bam.fofn"

python SQANTI-SIM/src/cage_sim.py train \
        --gtf $isoforms \
        --genome $genome \
        --CAGE_peak $CAGE \
        --SR_bam $SR_bam \
        --dir human_WTC11_PacBio_FLAIR_ssCAGE
```

#### 3 Simulation of SQANTI-SIM validation datasets

```
# full-sim NanoSim
python sqanti-sim.py full-sim sample \
    --gtf $annotation \
    --genome $genome \
    -d $val_nanosim_dir \
    --read_type cDNA \
    --ont --CAGE \
    --iso_complex \
    --trans_number 50000 --short_count 20000000 --falseCAGE_prop 0.8 \
    --ISM 1000 --NIC 1000 --NNC 1000 --Fusion 1000 --Antisense 1000 \
    --GG 1000 --GI 1000 --Intergenic 1000 \
    -k 8 --seed 3654

# full-sim PBSIM3
python sqanti-sim.py full-sim sample \
    --gtf $annotation \
    --genome $genome \
    -d $val_pbsim_dir \
    --pb --CAGE \
    --iso_complex \
    --trans_number 50000 --short_count 20000000 --falseCAGE_prop 0.8 \
    --ISM 1000 --NIC 1000 --NNC 1000 --Fusion 1000 --Antisense 1000 \
    --GG 1000 --GI 1000 --Intergenic 1000 \
    -k 8 --seed 3654
```

### 4 SQANTI-SIM benchmark isoform identification pipelines

#### 4.1 SQANTI-SIM simulation

```
genome="GRCh38.primary_assembly.genome.fa"
annotation="gencode.v43.annotation.gtf"
ont_dir="ont_sim_data"
pb_dir="pb_sim_data"

# full-sim NanoSim
python sqanti-sim.py full-sim sample \
    --gtf $annotation \
        --genome $genome \
        -d $ont_dir \
        --read_type cDNA \
        --ont --illumina --CAGE \
        --iso_complex --diff_exp 2 \
        --trans_number 50000 \
        --ISM 5000 --NIC 5000 --NNC 5000 \
        --long_count 20000000 \
        --short_count 60000000 \
        -k 8 --seed 3654

# full-sim PBSIM3
python sqanti-sim.py full-sim sample \
        --gtf $annotation \
        --genome $genome \
        -d $pb_dir \
        --pb --illumina --CAGE \
        --iso_complex --diff_exp 2 \
```

```
        --trans_number 50000 \
        --ISM 5000 --NIC 5000 --NNC 5000 \
        --long_count 4000000 \
        --short_count 60000000 \
        -k 8 --seed 3654
```

#### 4.2 Run analysis pipelines

##### 4.2.1 FLAIR

```
genome="GRCh38.primary_assembly.genome.fa"
reduced_annotation="sqanti-sim_modified.gtf"
lr="PBSIM3_simulated.fasta"

flair 123 \
        -r $lr \
        -g $genome \
        -f $reduced_annotation \
        -o $out_dir/flair_pb_LO \
        --temp_dir $out_dir/tmp_flair \
        --threads 8 \
        --check_splice
```

##### 4.2.2 FLAIR+Illumina

```
flair 123 \
        -r $lr \
        -g $genome \
        -f $reduced_annotation \
        -o $out_dir/flair_pb_LS \
        --temp_dir $out_dir/tmp_flair \
        --shortread $out_dir/Illumina_simulated.filtered.SJ.tab \
        --threads 8 \
        --check_splice
```

##### 4.2.3 TALON

```
# Mapping reads
minimap2 -ax splice:hq -uf --MD -t8 \
        $genome \
        $lr > $out_dir/PBSIM3_simulated.aln.sam

#Flagging reads for internal priming
talon_label_reads \
        --f $out_dir/PBSIM3_simulated.aln.sam \
        --g $genome \
        --t 8 \
        --o $out_dir/talon_pb_aln

# Initializing a TALON database
talon_initialize_database \
        --f $reduced_annotation \
        --g GRCh38 \
        --a gencode_v43 \
        --o $out_dir/talon_pb_db
```

```
# Running TALON
echo "pbsim,pbsim_pb,PacBio,$out_dir/talon_pb_aln_labeled.sam" > $out_dir/config.csv
talon \
        --f $out_dir/config.csv \
        --db $out_dir/talon_pb_db.db \
        --build GRCh38 \
        --threads 8 \
        --o $out_dir/talon_pb_run

# Filtering transcriptome for isoform-level analysis
talon_filter_transcripts \
        --db $out_dir/talon_pb_db.db \
        -a gencode_v43 \
        --o $out_dir/talon_pb.filtered_transcripts.csv

# Obtain custom GTF
talon_create_GTF \
        --db $out_dir/talon_pb_db.db \
        -b GRCh38 \
        -a gencode_v43 \
        --whitelist $out_dir/talon_pb.filtered_transcripts.csv \
        --o $out_dir/talon_pb.transcriptome
```

###### 4.2.4 IsoSeq

```
isoseq cluster2 \
        $out_dir/flnc.fofn \
        $out_dir/isoseq_pb_clustered.bam \
        --num-threads 8

pbmm2 align \
        $genome \
        $out_dir/isoseq_pb_clustered.bam \
        $out_dir/isoseq_pb_clustered.aligned.bam \
        --preset ISOSEQ \
        --sort \
        --num-threads 8

isoseq collapse \
    $out_dir/isoseq_pb_clustered.aligned.bam \
        $out_dir/isoseq_pb_clustered.gff \
        --do-not-collapse-extra-5exons
```

###### 4.2.5 IsoSeq+SQ3Rules

```
python sqanti3_filter.py rules \
    -j $filterjson \
    -o sq3filter -d $out_dir \
    --gtf $corrected_gtf $SQ3_class_file
```

##### 4.3 SQANTI-SIM evaluation

```
python sqanti-sim.py eval \
        --transcriptome $iso_dir/flair_ont_LO/flair_ont_LO.isoforms.gtf \
```

```
--gtf $reduced_gtf \
--genome $genome \
-o flair_ont_LO -d $out_dir/flair_ont_LO \
-i $trans_index \
-k 8 \
--min_support 5 \
--coverage Illumina_simulated_SJ.out.tab
```
